# Network and pathway expansion of genetic disease associations identifies successful drug targets

**DOI:** 10.1101/2020.04.22.051144

**Authors:** Aidan MacNamara, Nikolina Nakic, Ali Amin, Cong Guo, Karsten B. Sieber, Mark Hurle, Alex Gutteridge

## Abstract

It is widely accepted that genetic evidence of disease association acts as a sound basis for the selection of drug targets for complex common diseases and that propagation of genetic evidence through gene or protein interaction networks can accurately infer novel disease associations at genes for which no direct genetic evidence can be observed. However, an empirical test of the utility of combining these beliefs for drug discovery has been lacking.

In this study, we examine genetic associations arising from an analysis of 648 UK Biobank GWAS and evaluate whether targets identified as proxies of direct genetic hits are enriched for successful drug targets, as measured by historical clinical trial data.

We find that protein networks formed from specific functional linkages such as protein complexes and ligand-receptor pairs are suitable for even naïve guilt-by-association network propagation approaches. In addition, more sophisticated approaches applied to global protein-protein interaction networks and pathway databases, also successfully retrieve targets enriched for clinically successful drug targets. We conclude that network propagation of genetic evidence should be used for drug target identification.

## Introduction

A number of studies have shown empirically that genetic evidence provides a sound basis for the selection of new drug targets and the repurposing of existing drugs to new indications^1,2^. However, there are several reasons why individual genes might be missing direct genetic evidence associating them to diseases for which they could be used as drug targets. Therefore, various forms of network and pathway-based analyses have been proposed as a way to identify these ‘missing’ targets^3^ by integrating the results of genome-wide association studies (GWAS)^4^, gene interaction networks and signaling pathways^5,6^.

The hypothesis that some form of genetic association linking a gene to a disease makes the protein product of that gene a plausible drug target has a straightforward theoretical underpinning and reasonably strong empirical evidence to support it. The theoretical rationale is that genetic association, in contrast to most other forms of genomic association analysis, implies a clear causal relationship between changes in the activity of a gene product in humans and changes in the risk of developing the associated disease. This ability to confidently assign causation is due to the lack (in most common diseases outside cancers) of any plausible molecular mechanism for how the presence of disease could affect the DNA sequence and implies that pharmacological modulation could plausibly be expected to phenocopy the genotypic effect. To test this hypothesis, Nelson et al.^2^ showed through analysis of historic drug discovery programs, that genes with a direct genetic link to a disease have comprised 2% of preclinical drug discovery programs, compared to 8.2% of approved drugs. This implies that those targets with direct genetic evidence are more likely to succeed and therefore progress to approval than those without. Likewise, Cook et al.^1^ showed in an analysis of AstraZeneca’s drug discovery pipeline that projects in Phase II that had genetic evidence were successful 73% of the time compared to only 43% of the time for projects without genetic evidence.

These statistics raise an important question however: If genetics is a good way to select drug targets, why (to use the numbers cited by Nelson *et al*.) do 93.8% of approved drug targets not have direct genetic evidence linking them to the disease for which they are approved? The most likely answer to this is that, despite the exponential increase in the last few decades in our ability to genotype human subjects, our ability to measure genetic associations to the true disease phenotypes relevant for drug discovery is still limited, leading to ‘missing’ genetic associations. Most obviously, the majority of disease phenotypes for which GWAS are performed are related to risk of acquiring disease rather than progression or severity of symptoms of disease, which are usually (with notable exceptions such as cardiovascular disease^7^) the focus of current clinical practice and hence drug development. Even in those cases where the phenotype for which we have genotypic associations perfectly matches the phenotype of relevance for drug development, we may have limited power to detect genetic association due to the size of genotyped cohorts. Also, there may be an absence of suitable genetic instruments or we lack the ability to confidently map disease association signals to their *cis* or *trans* effector genes.

Where power to detect associations is an issue, one way proposed of detecting ‘missing’ genetic association is by using biological networks as a source of prior knowledge, as the propagation of genetic signals through those networks has been proposed as a ‘universal amplifier’^3^ that would improve our ability to find disease associated genes. Again, the theoretical rationale here is straightforward: As genes tend to interact with other genes that perform related cellular functions^8^, it should be possible to infer from the existence of a genetic disease association at one gene a link between that same disease and any other genes that interact with the original gene. Here we define these other genes as ‘proxy’ genes.

There are many approaches to defining proxy genes. Given a hypothetical model of a classical molecular signaling pathway (Figure 1A), consisting of ligand-receptor binding, protein complex formation, a kinase signaling pathway, and downstream nucleic effects (e.g. transcriptional regulation), we can define different functional categories to look for genes that interact with one or more disease-associated gene. The most conservative, but also naïve, approaches, simply look at a gene’s closest neighbors across the different functional categories. For example, if ligand *a* in Figure 1A is associated with a disease, an obvious strategy would be to look for potential drug targets in its binding receptors (Figure 1B-i). Other high-confidence functional interactions also make sense in this context, such as looking at stable protein complex partners of a disease-associated protein (*b* and Figure 1B-ii). Less conservative approaches might extend this strategy to first and second neighbors in the pathway of the disease-associated gene, or indeed extend the search to all genes in the pathway (gene *c* and Figure 1B-iii). More advanced algorithms try to infer an optimal subset of proteins to choose based on a combination of the patterns of direct genetic association and connections between proteins (e.g. algorithms such as Random-Walk that define disease-associated network modules based on these premises - genes *d* and *e*, and figure1B- iv).

**Figure 1:**
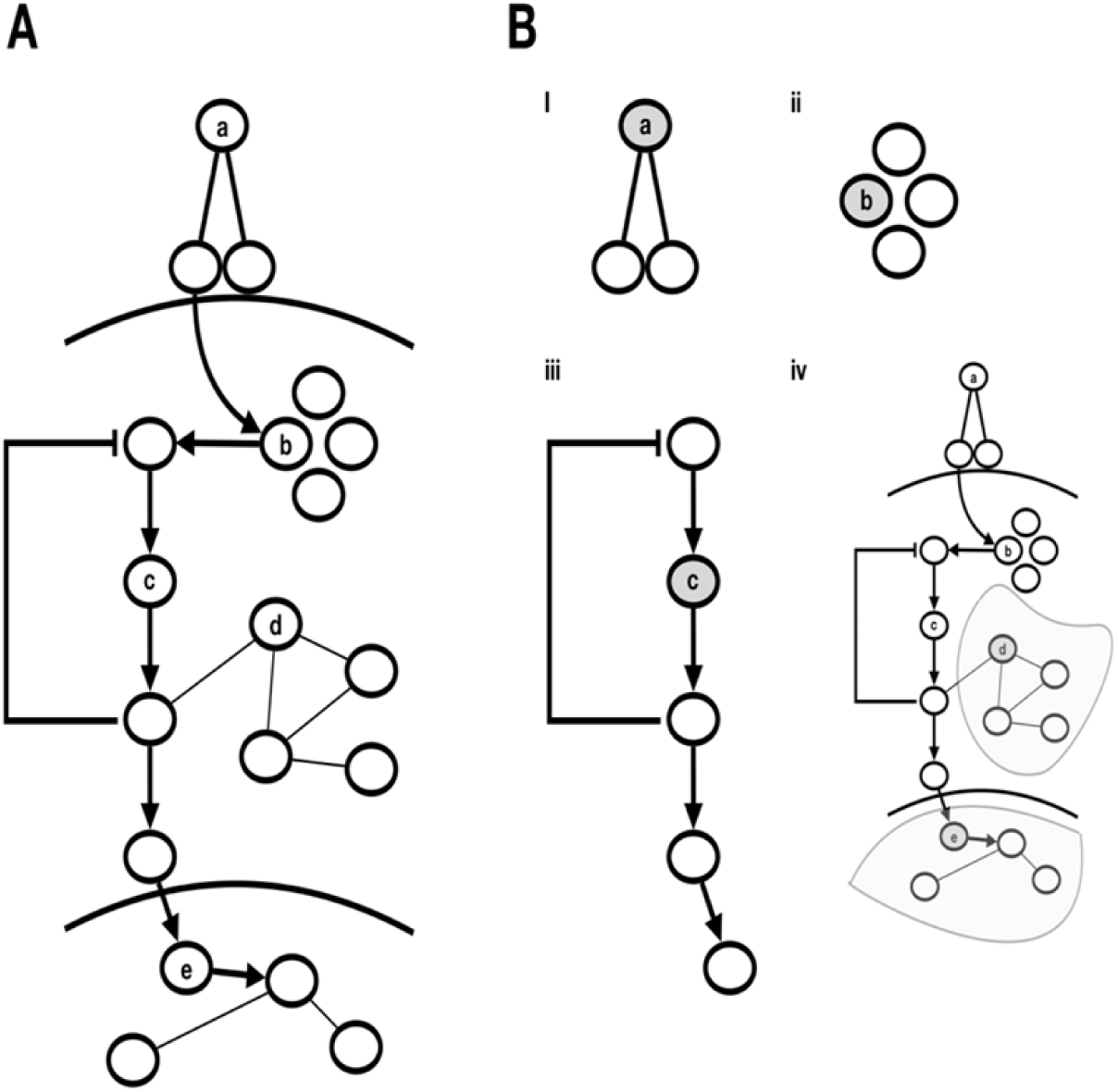
A schematic of a hypothetical model of a classical signalling pathway (panel A). *‘a’* is an extracellular ligand that binds to a multimeric receptor. *‘b’* is a member of a complex that triggers a pathway of protein kinases, of which *‘c’* is a member. *‘d’* is a possible regulator of that pathway but is not included in the canonical pathway definition (interactome resources would show *‘d’* interacting with members of the signalling pathway). *‘e’* is a transcription factor that regulates downstream expression. Panel B, i-iv show additional possible drug targets defined across functional categories, as detailed in the main text.

Several previous studies have applied the concepts of network propagation to disease association, but none have directly and systematically addressed the question of whether such approaches can maintain the empirically observed success of directly genetically associated genes when selecting drug targets, which is what we focus on in this study. A typical approach exemplified by Liu et al.^9^ takes the results of a set of GWAS (9 asthma related GWAS in this case), computes gene level scores and identifies a module or subnetwork of genes within a larger global PPI network that contains both known disease associated genes as well as a selection of novel targets. Nakka et al.^10^ take a similar approach using PEGASUS to compute gene scores and HotNet2^11^ to define the modules. Carlin et al.^12^ formulate a general scheme for performing these types of analyses and include infrastructure for storing and querying the derived networks in NDEx, a database of biological networks. Other approaches such as NetWAS^13^ derive new networks from molecular data that are then used alongside machine learning tools to produce systems that can re-rank GWAS output to prioritize genes with weak or even below threshold significance. Our own analysis of machine learning and network diffusion-based methods for inference of new disease associations through biological networks suggests that many of these methods^14^ perform very similarly and major differences in performance are driven more by the choice of the underlying biological network. Probably the largest systematic assessment of the use of network information to identify disease associations in complex diseases comes from the ‘Disease Module Identification DREAM Challenge’^15^. The inference task in this challenge is distinct from ours in that they aim to derive functional modules from networks without using disease association data directly. Instead disease association data is used to annotate and validate the function and biology of the derived networks. Another recent review has also benchmarked network algorithms using a different set of performance metrics and showed that network propagation performs well for target prediction^16^.

In this study, we first define a list of ‘high confidence genetic hits’ (HCGHs), which represent genes for which there is both a clear genetic association derived from GWAS and a clear mapping of the association to the gene through colocalization of the genetic disease association with an expression quantitative trait locus (eQTL). Then, we define genetic ‘proxy’ genes using various network and pathway analysis methods and sources of network prior knowledge. Finally, we measure the enrichment of successful drug targets for the given disease for both HCGHs and proxy genes with the aim of determining whether proxy genes are enriched for clinically successful targets and which methods are best suited for drug target selection.

## Methods

### GWAS data and the Definition of High-Confidence Genetic Hits (HCGHs)

UK Biobank (UKB) GWAS were selected for inclusion if a phenotypic match could be made between the (Medical Subject Headings) MeSH annotation of each trait and the MeSH annotations for indications with drug target success/failure data available from Citeline’s Pharmaprojects data (https://pharmaintelligence.informa.com/products-and-services/data-and-analysis/pharmaprojects, see Clinical Data section). This match was performed by fuzzy MeSH matching where one or more of the following conditions was true:

- If the relationship was a MeSH parent-child connection.
- Co-occurrence in literature abstracts significantly more often than random.
- Where at least one of two ontology-based methods which take into account the entire ontology structure^17,18^ gave a positive match.

For each GWAS, a set of genes were identified as ‘high confidence genetic hits’ (HCGHs) using colocalisation of the GWAS summary statistics with GTEx eQTLs. Colocalisation was performed^19^ followed by filtering such that colocalisation eGenes were selected to give 1 or 0 HCGHs for each disease-associated locus in the genome where:

- The eGene is protein coding *AND*
- The GWAS p-value ≤ 5e-8 *AND*
- The eQTL p-value ≤ 1e-4 *AND*
- The GWAS/eQTL colocalisation p12 ≥ 0.8 AND
- Where multiple such eGenes pass the above criteria for a single locus the eGene with the highest posterior probability of colocalisation (H4, p12) across all tissues was selected.

Only GWAS with ≥ 1 HCGH and ≥ 1 drug target with success/failure data available were retained in the analysis. Because of the fuzzy MeSH matching, some surgical GWAS traits were captured by this method but removed before further analysis. This resulted in 648 GWAS covering 170 individual MeSH traits linked to 14374 distinct HCGH-GWAS combinations, and 1045 distinct drug targets with success/failure data. All methods were subsequently tested on these traits, genetic hits, and drug targets using the full protein-coding gene list as the background gene universe (22758 genes).

### Clinical Data

We extracted data from Citeline’s Pharmaprojects database (https://pharmaintelligence.informa.com/products-and-services/data-and-analysis/pharmaprojects, downloaded 5^th^ August 2017), reformatting available XML data into a single tab-delimited form having one row for each asset. Each asset may be linked to one or more targets, whether due to specific action at a complex or non-specific action against multiple targets. Each asset may also be linked to progression against one or more indications, each with its own pipeline status.

We classified the 116,532 asset-indication pairs into one of 3 categories: ‘Succeeded’, ‘Failed’ or ‘In Progress’, based on the status listed in Pharmaprojects for each indication. The 9,026 in the ‘Succeeded’ category consists entirely of ‘Launched’ pairs. The 79,824 asset-indication pairs with ‘Failed’ status consist of ‘Discontinued’ (24%), ‘No Development Reported’ (76%), ‘Withdrawn’ (<1%) or ‘Suspended’ (<1%) asset-indication pairs, while the remaining 27,295 pairs, which typically list the individual clinical or preclinical phase, are classified as ‘In Progress’.

We then classified the failures. Based on a collation of data from several text fields in Pharmaprojects (Key Event Detail, Overview, Phase III, Phase II, etc.), we manually deduced the pipeline status (Preclinical, Phase I, Phase II, Phase III) of each indication and from ‘Key Event History’, the date of failure for the ‘Failed’ asset-indication pairs where available. In general, assets with a single indication were straightforward to assign based on the clinical phases that were mentioned; for those with multiple indications, we looked for phrases which linked a specific indication with a specific clinical phase. We did not include instances when a clinical phase did not actually appear to be undertaken based on the available text, such as if the trial was ‘planned’ or ‘under consideration’. 26% of the failures reported in Pharmaprojects could be determined to be clinical failures by this method.

To group similar findings together and prepare them for matching to evidence types, we assigned each of the 1,340 unique indications in Pharmaprojects to one of 1,063 Medical Subject Heading (MeSH) disease terms. 2,588 asset-indication pairs with indications classified as ‘Ideopathic disease, unspecified’, ‘Not applicable’, ‘Undisclosed’ and ‘Unspecified’ or any of the 15 diagnosis terms were not mapped and were not processed further, as a successful marker of the disease is not an indication that the disease has been therapeutically treated. We also used Pharmaprojects mappings for assets to human EntrezGene IDs to generate a list of 39,661 human target-asset pairs, correcting the single EntrezGene ID listed in Pharmaprojects which is not currently used (SCN2A, from 6325 to the correct 6326). We then produced a list of asset-EntrezGene-MeSH combinations, indicating whether the asset binds to a single target or multiple targets, and whether it is being progressed against a single indication or multiple indications.

We then grouped these 80,804 asset-target-indication triples (that is, those asset-indication pairs with a human target) into 27,064 unique target-indication pairs, noting which of these assets were labelled as interacting with one target (‘Selective’), and those which interacted with more than one target (‘Non-Selective’). Non-Selective assets could represent poly- pharmacology or binding of the asset to a complex of targets. If at least one ‘Selective’ asset for a given target-indication pair was identified as successful, then the target-indication pair was classified as ‘Succeeded’. Of the remaining target-indication pairs, if at least one ‘Selective’ asset had a clinical failure then it was classified as ‘Clinical Failure’. We then processed the data in the same way for ‘Non-Selective’ assets. The remaining data were processed in the same order for ‘Preclinical Failures’. Those target-indication pairs which had not yet been indicated as failures or successes were then identified as ‘In Progress’, in that no record of success or failure yet exists in Pharmaprojects for these target-indication pairs. For each pair, we also recorded the furthest clinical phase achieved by any past or current asset.

For this analysis, we utilized those target-indication pairs classified as ‘Succeeded’ as our positive set, and those classified as ‘Clinical Failure’ as our negative set.

### Evaluation of Methods for Proxy Gene Set Definition

The following methods were used to define proxy gene sets for the HCGHs for each GWAS. See below for details on data sources used:

- Complex: All genes sharing a protein complex with a HGCH
- Ligand Receptor: All genes in a ligand-receptor pair with a HCGH
- Network First Neighbor: All first-degree interactors of a HCGH
- Network Second Neighbor: All first and second-degree interactors of a HCGH
- Pathway: All genes in the same pathway as a HCGH
- Pathway First Neighbor: All first-degree interactors that also share a pathway with a HCGH
- Pathway Second Neighbor: All first and second-degree interactors that also share a pathway with a HCGH
- Random: 10000 randomly selected protein coding genes from the background of 22758
- Hotnet*: All genes found within a HotNet2 network module (see below)
- Pascal/MAGMA identified genes (see below)

Complex: Data downloaded from https://www.ebi.ac.uk/complexportal/home on 21/01/2019. Ligand-Receptor: Data sourced from Ramilowski *et al*^20^ and parsed from Metabase (https://portal.genego.com/). In brief, Metabase was parsed for ligand-receptor related keywords in the interaction metadata. Non-specific interaction types were then removed. Networks: Multiple different gene networks were used as follows:

- OmniPath: The OmniPath interaction file was downloaded 14/02/2019
- STRING: Human interaction data was downloaded from STRING on 14/02/2019
- HuRI: Data downloaded from http://interactome.baderlab.org/
- InBio Map: Data downloaded from https://www.intomics.com/inbio/map.html#downloads

For defining first and second neighbours from the network sources, each network was converted into an iGraph object in R. The iGraph functions neighbors() and neighborhood() were then used to find first/second neighbours, respectively. The list of first and second pathway interactors within pathways was created using Metabase pathway maps. First interactors for a gene were defined as all upstream and downstream direct interactors across all pathways. Then, the process was repeated starting with the first interactors, thus creating a list of second interactors.

### Enrichment Calculations

The enrichment of successful drug targets within the HCGHs and proxy gene sets was calculated for each GWAS/method pair. For each pair a 2×2 contingency table was constructed as follows:

|  | Success | Failure |
| --- | --- | --- |
| Hit | *a* | *b* |
| Not Hit | *c* | *d* |

Depending on the method, some tables had *a* and/or *b* equal to 0 (i.e. no overlap between hits and failed or successful drugs). Tables with both *a* **and** *b* equal to 0 were removed. Where *a* **or** *b* were equal to 0, 0.5 was added to each cell in the contingency table (Haldane Correction^19^). The reason for this was to preserve information where otherwise the odds ratio would be undefined or infinite. To calculate an odds ratio and significance of enrichment for each method, a stratified Fisher’s Test was then used (the Cochran–Mantel– Haenszel test), across all GWAS for each method. Odds ratios and 80% confidence intervals were then reported to measure by-method enrichment of successful drug targets.

### Network Propagation

HotNet2^11^ was used to define HCGH-enriched network modules. For this method, genes found in network modules, excluding the seed HCGHs, were defined as hits. HotNet2 takes 2 inputs, a network and a gene list that defines the seed genes (in our case, the HCGHs) and their associated genetic scores. For each GWAS/network combination, the HCGH gene set was used as the input gene list and the score for each HCGH was derived from the p12 colocalisation probability for that gene. The p12 probabilities were transformed by – log(1-p12, base=2). For the purpose of this study, the consensus modules were used and all genes contained within these modules were defined as hits for the GWAS/network.

### Gene score and pathway enrichment calculation

Gene scores were calculated using two different algorithms: Pascal^21^ and MAGMA^22^. Pascal was run with default settings, using the ‘sum’ gene scoring method. The ‘empirical’ pathway enrichment p-value was taken as the measurement of pathway enrichment. For both Pascal and MAGMA the 1KG LD matrix was used and the definition of the gene locus was the gene body +/- 50kb. A number of different gene-sets were used as input for both methods: 1) Metabase pathway maps, 2) Reactome pathways, 3) <u>DREAM</u> networks consensus PPI modules, 4) <u>DREAM</u> networks consensus co-expression modules. Gene-set enrichment p-values were adjusted for multiple hypothesis testing using the BH method, calling pathways with the adjusted p-value < 0.05 significantly enriched for the tested GWAS trait. A manually curated list of HLA genes was excluded from both gene-set level analyses. We found that Pascal significantly outperformed Magma (supplementary Figure 6). Hence, we removed Magma from further analysis.

## Results

### Naïve Approaches

Our first approach is to look at a set of relatively naïve network expansion methods, the results of which are shown in Figure 2. For these methods the algorithm is simply the selection of first or first and second neighbors within the relevant protein-protein interaction network. Our positive control is the list of HCGHs for which there is clear, direct genetic association to disease. Consistent with previous work we confirm that such targets are significantly enriched for those which have proved to be successful (OR: 3.8; p < 1×10^−6^). Our negative control is a set of randomly chosen genes from the background set which we confirm to have no significant enrichment for successful drug targets (OR: 1; p = 0.8).

**Figure 2:**
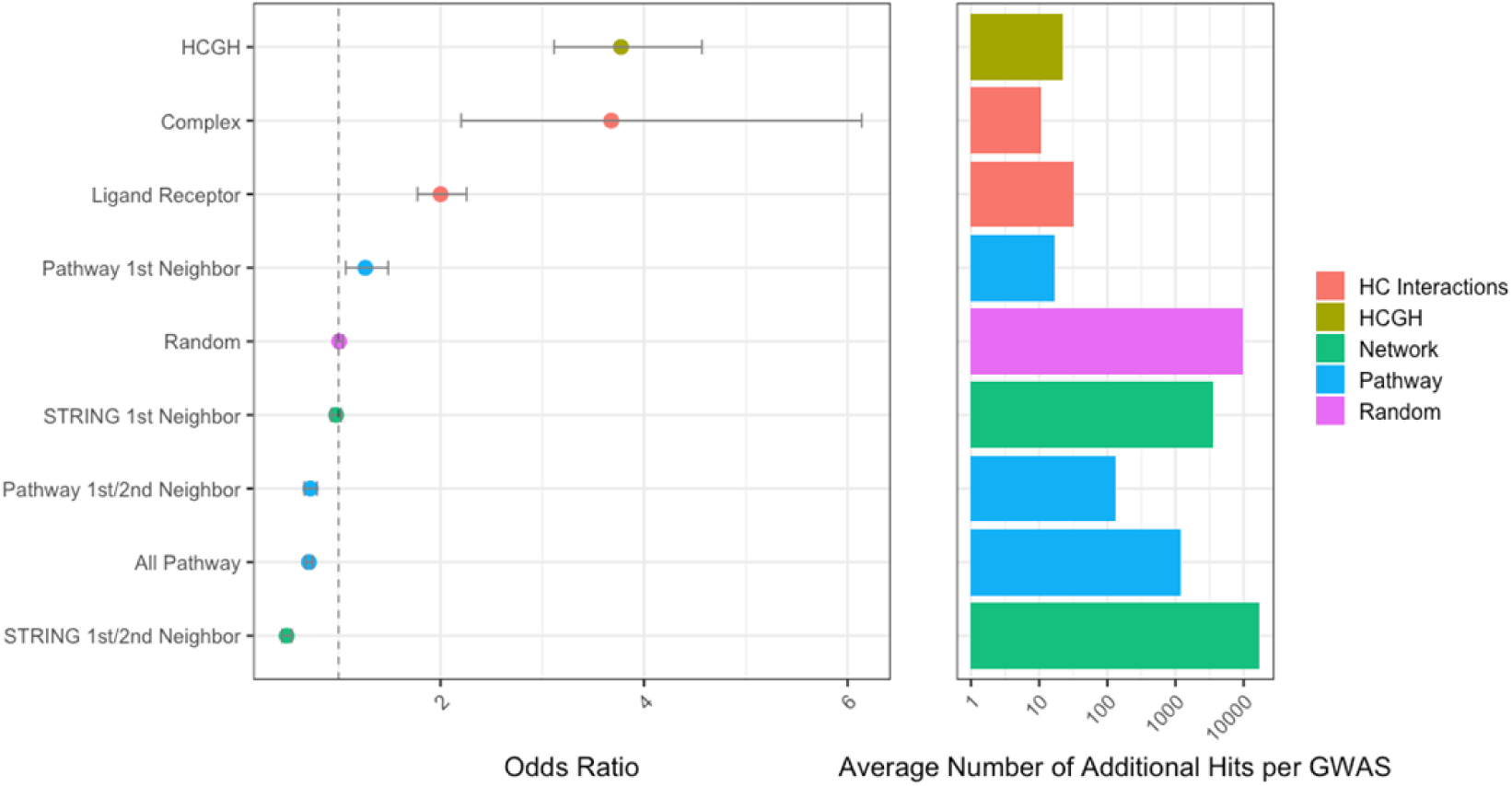
Enrichment of successful drug targets (left) and number of targets implicated (right) within HCGHs and additional target sets identified by network expansion using HCGH seeds. The colours refer to the interaction categories used for the expansion (red; high-confidence interactions – complexes and ligand-receptor pairs, green; protein-protein interaction network STRING, blue; interactions from pathways defined in Metabase)

The first network we examine comprises stable protein complexes. In this network we model each complex as a fully connected clique (i.e. every member of the complex is a first neighbor of every other member). Taking HCGHs and performing network expansion using this network adds ∼10 novel potential target genes to the average GWAS and those genes are enriched with successful drug targets to a similar level as the positive control (OR: 3.7; p = 1.4×10^−3^). This enrichment calculation (and all following calculations) is performed on the new proxy genes *only* with the original seed HCGHs removed. Since protein complexes comprise highly curated sets of genes that should have very high levels of shared cellular function, the result of observing high enrichment is not surprising, but it does confirm that this conservative level of network expansion is advisable in a target identification exercise.

The second network we examine comprises ligand-receptor pairs. In this network (which is not a simple 1:1 mapping), we model each ligand as being connected to all the proteins that comprise its receptors and vice-versa we connect each receptor subunit to all its possible ligands. Note that in this analysis we only consider first neighbors. We do not expand to second neighbors, which would have the effect of propagating genetic evidence from a ligand to its receptor and hence to *all* of that receptor’s ligands. Again, we find that the additional targets identified through this approach are enriched for successful drug targets (OR: 2; p < 1×10^−5^) and confirm that network expansion using this class of network is reasonable to perform when undertaking target identification.

The third network we use is STRING for which we measure success enrichment amongst first neighbors and the union of first and second neighbors of the HCGHs. We observe no enrichment for successful drug targets amongst first neighbors of the HCGHs (OR: 1; p = 0.5) and a significant enrichment of *failed* targets amongst the first and second neighbors of the HCGHs (OR: 0.5; p < 1×10^−5^). This second observation is worthy of comment as the apparent conclusion – that second neighbors of genes genetically associated to a given disease are significantly more likely than a random gene to *fail* as drug targets for that disease – is not intuitive. The reason we arrive at this conclusion comes from a property of the network and the way in which historically tested drug targets are distributed within it; namely that a small number of genes are very highly connected within the network (expected due to the scale-free topology of most biological networks) and that these genes happen to have been the focus of historical drug discovery efforts, which mean they have been tested in a high number of trials and that those trials contain a high proportion of failures. This effect is shown graphically in supplementary Figure 2. In both cases (first and first & second neighbors), the number of additional targets implied by network expansion is very large (1000s and even 10,000s of additional targets for most GWAS). The use of alternative networks to STRING can somewhat ameliorate the effect observed of enrichment of *failed* targets within first and second neighbors. However, in no network do such simple algorithms provide value in terms of target selection (supplementary Figure 1).

The fourth network is based on pathway maps taken from Metabase. In our first naïve analysis we consider a network where every pathway map is modelled as a clique – every member of the pathway connects to every other (Figure 1iv). Our other analyses take the pathway connectivity defined in Metabase pathways into account and restrict the expansion to first or first and second neighbors. As with the STRING network, taking the clique (OR: 0.7; p < 1×10^−6^) and first and second neighbor (OR 0.7; p < 1×10^−6^) approaches within Metabase pathways leads to an enrichment of failed drug targets for the same reasons as above. Taking first neighbors within the pathway does provide a small enrichment of successful drug targets (OR 1.26; p = 0.07) and a similarly small number of additional targets.

### Advanced Approaches

All methods used in the above analyses (naïve) rely on careful selection of highly curated protein interaction networks followed by the application of very simple – essentially trivial - algorithms to select first or first and second neighbors of the HCGH seed genes.

Unsurprisingly these algorithms perform very poorly when applied naively to a densely connected network such as STRING. An obvious and frequently used extension to these algorithms is to apply some form of network propagation. Here we use the HotNet2^10,11^ algorithm and search for enrichment of successful drug targets on four different protein interaction networks, as shown in Figure 3.

**Figure 3:**
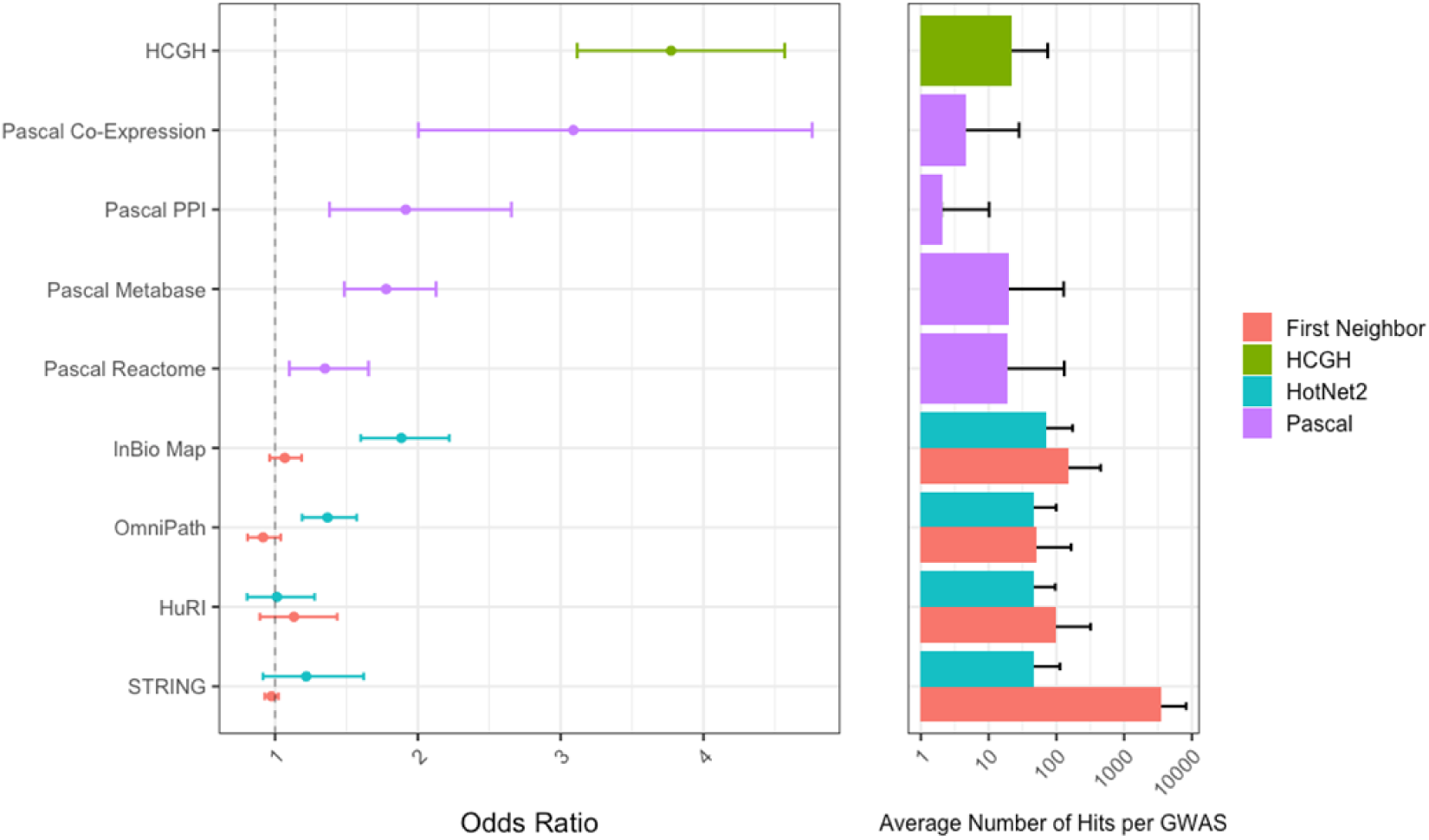
Enrichment of successful drug targets within HCGHs and proxy gene sets (left) and the number of additional potential targets identified (right). The different enrichment categories are colour-labelled; for network propagation, the enrichment performance of running HotNet2 is shown together with the enrichment gained from including first neighbours of HCGHs.

The genes found within modules detected by HotNet2’s network propagation and module selection algorithms (Figure 3; green) are significantly enriched for successful drug targets in the InBio Map and OmniPath networks (OR: 1.88/1.37; p < 1×10^−4^ / p = 3×10^−4^). HotNet2 does not reach significance with STRING (OR: 1.2; p = 0.42). The odds ratio point estimate for enrichment for HotNet2 applied to HuRI is also insignificant and close to unity, though with considerable error bars (OR: 1; p = 0.94). In all cases HotNet2 identifies 60-70 new targets through inclusion in the modules detected.

The final scenario we test is based on the pathway enrichment of gene scores that are derived from Pascal. We test what happens if we select as targets sets of genes that are both within a pathway or a network module that is itself significantly enriched for genetic association to a disease as measured by a GWAS (based on a Pascal gene score threshold) and have a nominally significant (P < 0.05) Pascal gene score to the same disease in the same GWAS. We use Pascal^22^ to test the performance of this strategy, which is also shown in Figure 3.

The genes found within pathways and modules detected by the Pascal algorithm (Figure 3; purple) are significantly enriched for successful drug targets across all tested sources of pathway gene sets and network modules, apart from the Reactome pathways. Pascal analysis on DREAM co-expression modules resulted in an enrichment close to that of HCGHs themselves with an OR: 3.09 and p = 3×10^−4^. Analysis of network modules, both PPI and co-expression variants, however, yielded a limited number of new targets (3-7), while the analysis of pathways yielded ∼30 new targets.

The full results across all methods can be found in supplementary Table 1 and supplementary Figure 3.

One possible reason that we observe drug targets with no genetic evidence is that our ability to find genetic associations between these targets and their respective diseases is hampered by underpowered association studies. This would suggest that our proxy targets should have some higher level of genetic association signal compared to random genes even if the signal does not reach genome wide significance. Figure 4 shows the distribution of gene level disease association scores calculated using Pascal^15^ for the proxy genes identified by each of the methods described above. Gene scores are given only for the GWAS trait implicated by the original seed HCGH. HCGHs themselves have consistently high (on a - log(P) scale) gene scores as one would expect (some HCGHs do not have significant gene scores calculated by Pascal as the colocalization used to define them can be driven by enhancers outside the gene body window used by Pascal). What is more revealing is that all the proxy gene sets identified in these networks have an average gene score higher than a random distribution and that the size of this effect largely tracks the enrichments observed above: unsurprisingly the effect is larger in the more advanced methods such as Pascal that use the genetic signal directly (Figure 4; right) but is also demonstrated for naïve methods (Figure 4; left, and supplementary Figure 4), and is true even based on very different underlying network structures (supplementary Figure 5). This observation is consistent with previous work that has shown that genes with nominally significant Pascal scores from a GWAS can be used with network information to predict genetic associations subsequently found in independent genetic studies for the same trait^14^.

**Figure 4:**
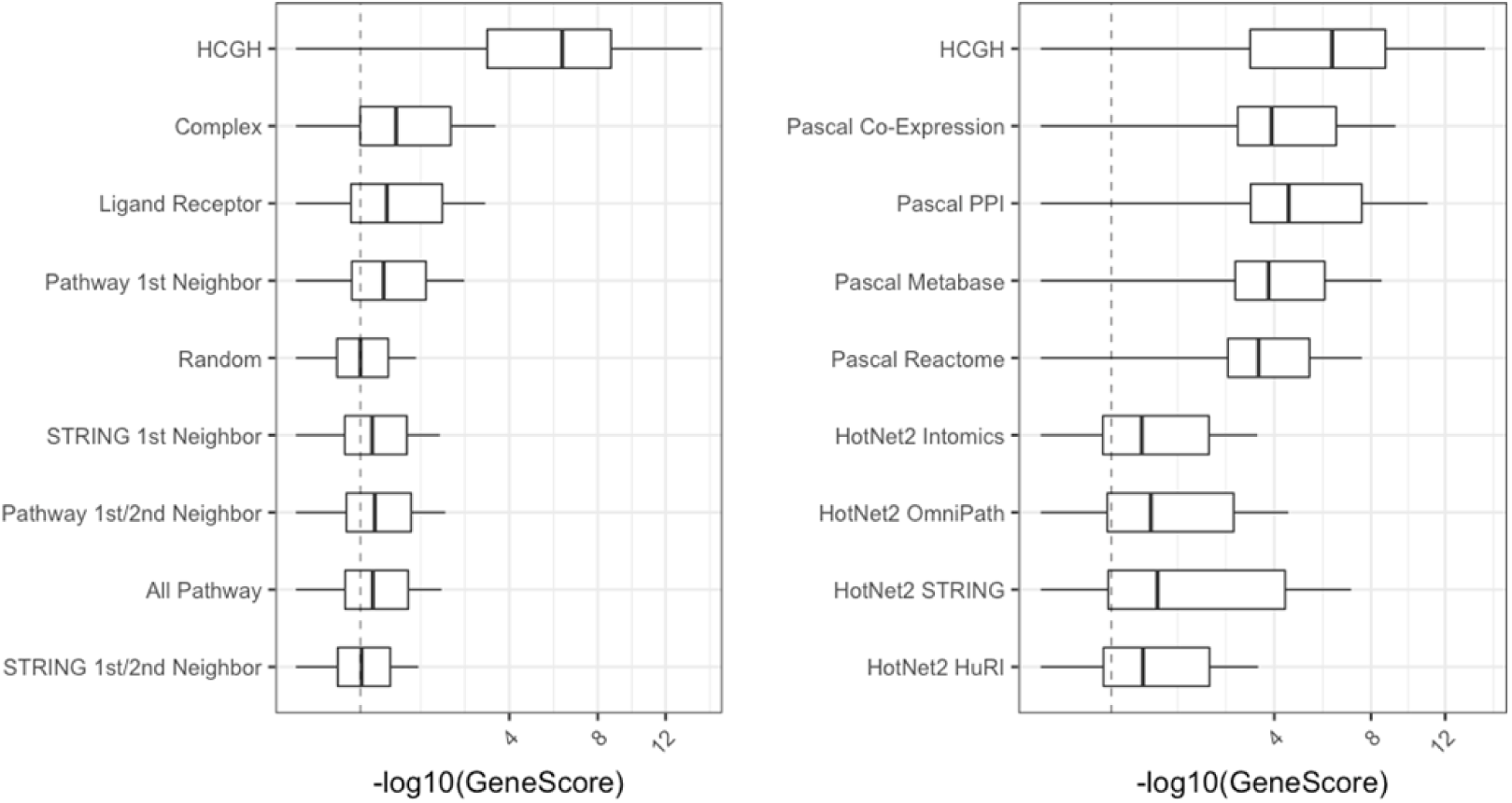
Pascal gene scores for HCGHs and all proxy genes found using each method indicated. Scores for the original seed HCGHs are excluded from the results across all the network expansion methods. The order of methods is the same as Figure 2 (left) and Figure 3 (right).

**Figure 5:**
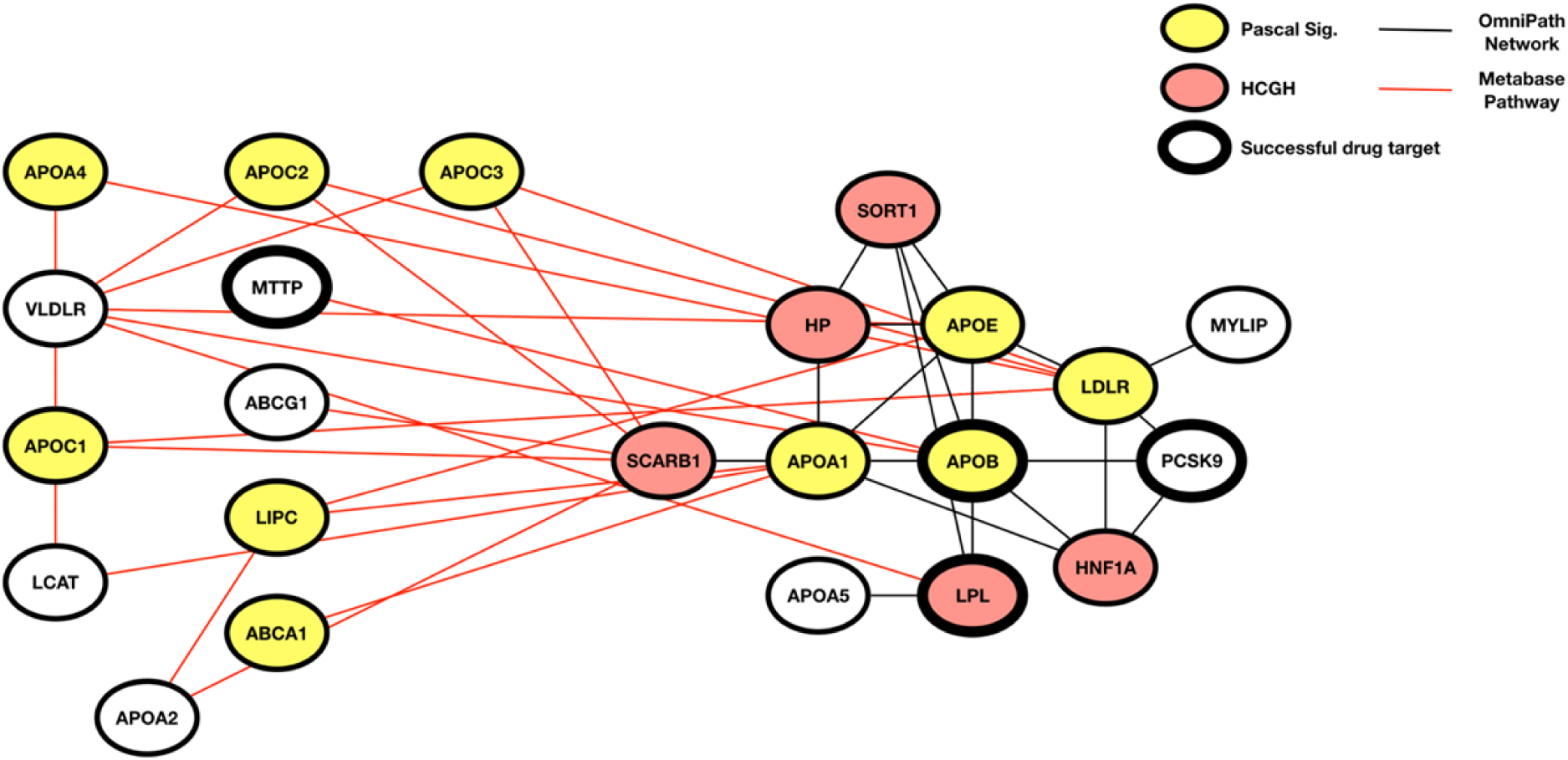
An example of where Pascal pathway enrichment and HotNet2 home in on the same pathway causal of Hyperlipidaemia. The Hotnet2 module (black interactions) was detected using HCGHs (red) from a Hyperlipidaemia GWAS and the OmniPath network. This module is enriched for 3 successful drug targets (bold node border), one of which, PCSK9, was not categorised as a HCGH in the input. The Pascal enriched pathway (red interactions) is the lipoprotein metabolism pathway from Metabase. Genes that have a significant Pascal gene score are highlighted in yellow: it can also be seen that an additional novel drug target was recovered (MTTP) using this method that did not have any type of genetic evidence associated with it.

## Discussion

Our results confirm the widely held intuition that inference of disease associations through a combination of direct causal evidence at single genes (provided by genetics) and propagation of that evidence through a protein interaction network (that captures genuine functional relationships) is a sound strategy for drug discovery. We go further than this though in providing a more thorough empirical understanding of the types of protein networks for which this strategy is valid and the types of algorithms which should be used for propagation. We also provide additional quantitative understanding of the ways in which diffusion of disease association within a protein network is manifested in observable genetic associations.

Our headline conclusion for simple first or first and second neighbor ‘guilt-by-association’ approaches to target identification is that these are valid and useful for networks of protein complex members or ligand receptor pairs, but not for other commonly used forms of network or pathway information. An open question we do not answer is whether other specific interaction types exist that would have similar properties to complexes or ligand- receptor pairs. Our observation of weak but significant enrichment of successful drug targets amongst first neighbors within pathway maps and an enrichment of weak genetic associations within HCGH PPI first neighbors may well imply that such networks do exist. Kinase-substrate or phosphatase-substrate networks would be obvious choices to inspect in that they often define the core elements of signaling pathways. Alternatively, enzymatic pathways (linking enzymatic producers of a compound to consumers) could also be tested especially where metabolomic QTL or other evidence exists for associating the cognate metabolites to disease as well^23^.

Our second conclusion is that more advanced network propagation algorithms can provide the ability to detect patterns of useful disease association within even densely connected proteome-scale interaction networks such as InBio Map and genome-scale signaling pathway maps such as OmniPath. This effect is primarily due to the ability of HotNet2 to exclude as potential targets large numbers of genes that are close to HCGH seeds, but do not sit within a coherent pattern of disease association within the network. A weakness of our study is that we do not test other network propagation methods. However, many such methods are based around some version of the random walk with restart algorithm or a mathematically equivalent conception and in previous work we have showed that many such algorithms perform equivalently on a highly related problem^14^. One potential avenue for development in this area would be in graph based deep learning that could explicitly model other additional sources of disease association such as those from target information integration platforms such as Open Targets^24^. Figure 5 also highlights the importance of these more advanced approaches in discovering the mechanisms behind genetic association with disease. Here we have two independent methods, Pascal and HotNet2, using 2 different network sources (Metabase pathways and OmniPath), homing in on the same biology that underlies hyperlipidaemia.

Our final conclusion is that what these processes are modelling is the diffusion of disease association. Causal disease association is the property one fundamentally looks for in drug targets and genetic association is one of, if not the, best way to detect such associations.

The first limitation of our study to recognize is that we only test network propagation of genetic evidence and in fact restrict ourselves to one specific form of genetic evidence, namely colocalization of eQTL and disease association loci. However, given the evidence supporting truth set enrichment from colocalization, we anticipate that it would have a relatively low false positive rate for identifying truly disease associated genes. The thresholds we use mean that the evidence for disease association itself at a given locus should be robust as well as the evidence for colocalization of the disease locus with gene eQTLs. The major source of false positives will be through loci containing either pleiotropic eQTL signals or many independent eQTLs leading to misassignment of the effector gene. The downside of this approach however is that we also have a high false negative rate in that there will be many genes with strong and obvious genetic evidence for a trait that we miss (protein coding variants for instance). Our aim however is not to perfectly catalog all genetically associated genes for these traits (this is left as an exercise for the reader), but rather to test the validity of our network and propagation models given some reliable form of genetic evidence. Our expectation would be that the same approaches would be valid no matter the source of the genetic association evidence, whether it be eQTL based or from protein coding variants or even based on rare Mendelian genetics; though we have not formally shown this.

The more important limitation of our study arises from the way in which we measure the performance of the various methods and networks using historical drug discovery data. The limitation of this data is that it is highly biased and has a large amount of missing data in terms of providing a true measurement of the universe of good drug targets for a given disease. Both effects are well known and described; firstly, genes are not chosen as drug targets in an unbiased way; instead certain families of genes (G-protein coupled receptors and protein kinases for instance) are much more likely to be chosen as targets^25^ compared to others. This is because of properties, such as druggability, that are entirely orthogonal to the strength of disease association alone. Also, genes that themselves have been highly studied in terms of their molecular and cellular function are more likely to be chosen as targets compared with genes of unknown or poorly understood function. In addition, targets that have been tested against a large number of diseases are more likely to have a higher proportion of failures than those which have only been tested against only a few diseases (supplementary Figure 2). This probably reflects the decreasing marginal cost of each additional clinical trial for a given drug since most of the typical preclinical and Phase I costs are already sunk. This in turn makes increasingly riskier trials for additional indications, based on weaker disease association evidence, worthwhile from a commercial risk-reward perspective. Secondly, the large amount of missing data arises simply from the fact that drug discovery activities and clinical trials especially are expensive and therefore relatively few of the potential targets for a given disease have ever been tested clinically.

It is important to bear in mind therefore that what we are measuring when looking at historical trial outcomes is not an unbiased measure of any given gene’s true *disease* associations, but rather a view on how useful a given evidence source or analytical method has been for choosing *drug* targets based on current and historical drug discovery practices. Dramatic changes in these practices in the future could render some of our conclusions obsolete, though the fundamental observation that genetic association itself is retained in molecular networks will remain valid. It is especially important to bear these facts in mind when considering our apparently counter-intuitive result that first and second neighbors of HCGHs for a given disease are enriched for *failed* drug targets against that disease. Taken naively that would imply that one should deliberately ignore potential drug targets that are first or second neighbors of HCGHs in a target identification exercise, but this would be a very odd conclusion that is hard to rationalize biologically. More realistically we would suggest that the true conclusion to draw from this part of our study is that such naïve approaches are not detectably better than a random selection of drug targets and that further work on the development of graph-based machine learning algorithms for the selection of drug targets based on genetics and other disease association information is therefore warranted.

## Acknowledgements

Many thanks to all our past and current colleagues within the GSK Human Genetics and Functional Genomics groups for their comments and feedback on this manuscript and the original research program. Special thanks to our colleagues in GSK Research Statistics for fruitful discussions on the statistical approaches used.

## Author contributions

A.M. and A.A. performed the network propagation and enrichment analyses. N.N. performed the gene and pathway level genetic enrichment analyses. K.G. and K.S. performed GWAS and colocalization analyses. M.H. performed the curation and analysis of clinical trial data. A.G., A.M. and N.N. wrote the manuscript. All authors contributed to the conception of the study and the development of the methods.

