## Supplementary Materials for "Network and pathway expansion of genetic disease associations identifies successful drug targets"

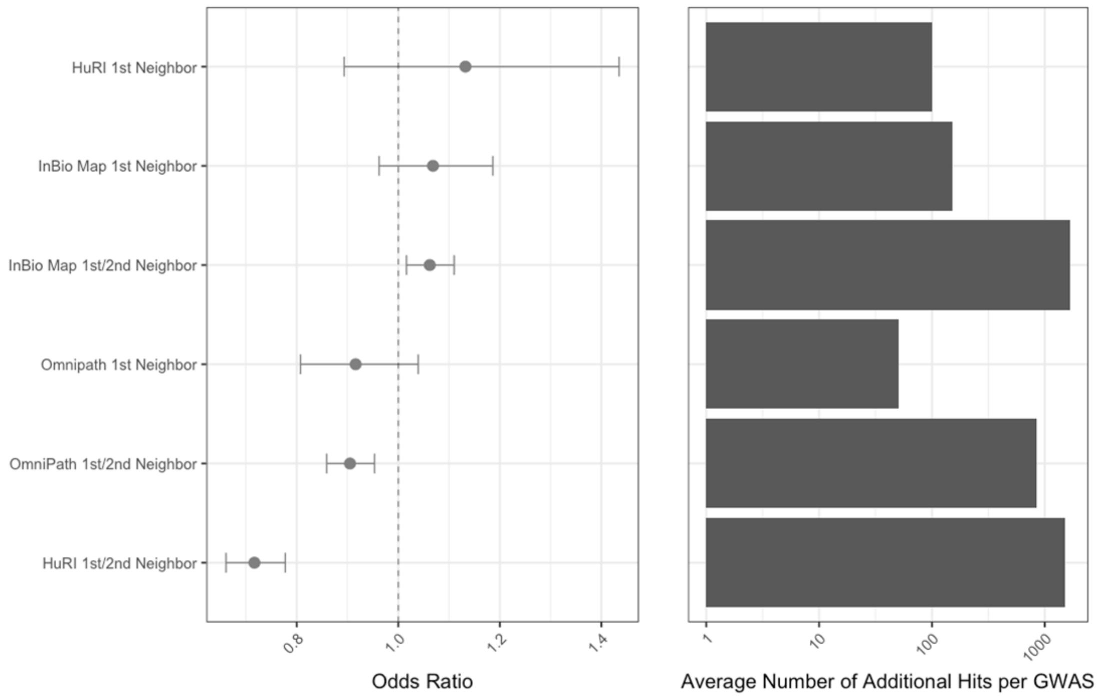


Supplementary Figure 1: Enrichment of successful drug targets (left) and number of targets implicated (right) using the naïve approach of first and second neighbors across different network sources (HuRI, InBio Map, STRING, OmniPath)

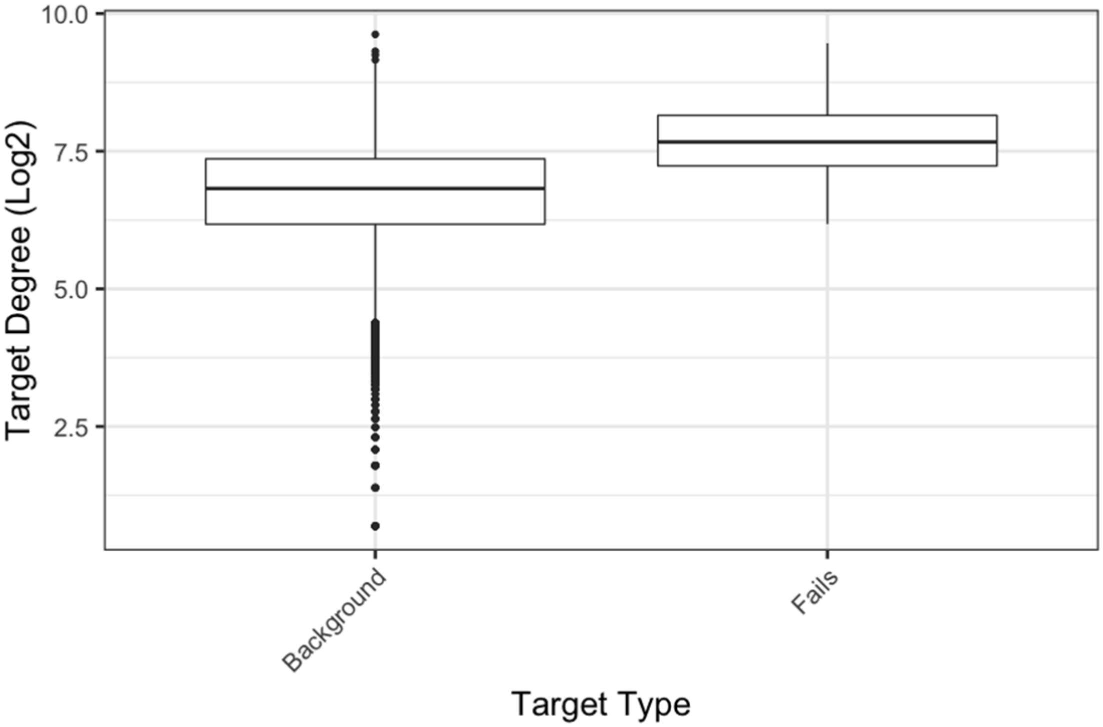


Supplementary Figure 2: A comparison of the node degree (the number of connections in a network that a gene has) between genes that have the most clinical failures *(n = 500)* against all other genes.

| Method | Odds | Confidence Intervals 80% | | P |
|---|---|---|---|---|
| HCGH | 3.77 | 3.12 | 4.57 | 0.0000 |
| Complex | 3.68 | 2.20 | 6.14 | 0.0014 |
| Pascal Co-Expresssion | 3.09 | 2.00 | 4.76 | 0.0003 |
| Ligand Receptor | 2.00 | 1.77 | 2.26 | 0.0000 |
| Pascal PPI | 1.92 | 1.38 | 2.66 | 0.0088 |
| HotNet2 Inbio Map | 1.88 | 1.60 | 2.22 | 0.0000 |
| Pascal Metabase | 1.78 | 1.48 | 2.13 | 0.0000 |
| HotNet2 OmniPath | 1.37 | 1.19 | 1.57 | 0.0035 |
| Pascal Reactome | 1.35 | 1.10 | 1.65 | 0.0587 |
| Pathway 1st Neighbor | 1.26 | 1.07 | 1.49 | 0.0728 |
| HotNet2 STRING | 1.22 | 0.92 | 1.62 | 0.4206 |
| HuRI 1st Neighbor | 1.13 | 0.89 | 1.43 | 0.5248 |
| InBio Map 1st Neighbor | 1.07 | 0.96 | 1.19 | 0.4200 |
| InBio Map 1st/2nd Neighbor | 1.06 | 1.02 | 1.11 | 0.0791 |
| HotNet2 HuRI | 1.01 | 0.80 | 1.28 | 0.9386 |
| Random | 1.01 | 0.97 | 1.04 | 0.8275 |
| STRING 1st Neighbor | 0.98 | 0.93 | 1.02 | 0.5227 |
| OmniPath 1st Neighbor | 0.92 | 0.81 | 1.04 | 0.3888 |
| OmniPath 1st/2nd Neighbor | 0.90 | 0.86 | 0.95 | 0.0150 |
| Pathway 1st/2nd Neighbor | 0.72 | 0.66 | 0.79 | 0.0000 |
| HuRI 1st/2nd Neighbor | 0.72 | 0.66 | 0.78 | 0.0000 |
| All Pathway | 0.71 | 0.68 | 0.74 | 0.0000 |
| STRING 1st/2nd Neighbor | 0.49 | 0.45 | 0.54 | 0.0000 |

Supplementary Table 1: The odds ratio and 80% confidence intervals of successful drug enrichment across all methods

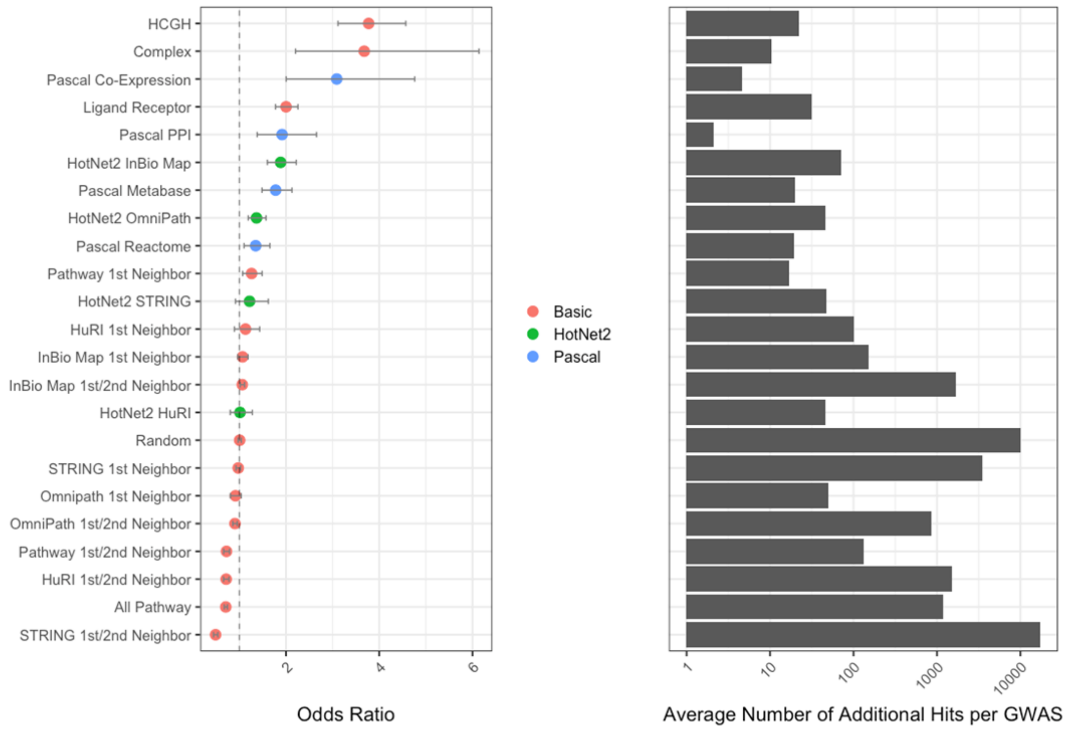


Supplementary Figure 3: Enrichment of successful drug targets (left) and number of targets implicated (right) across all methods.

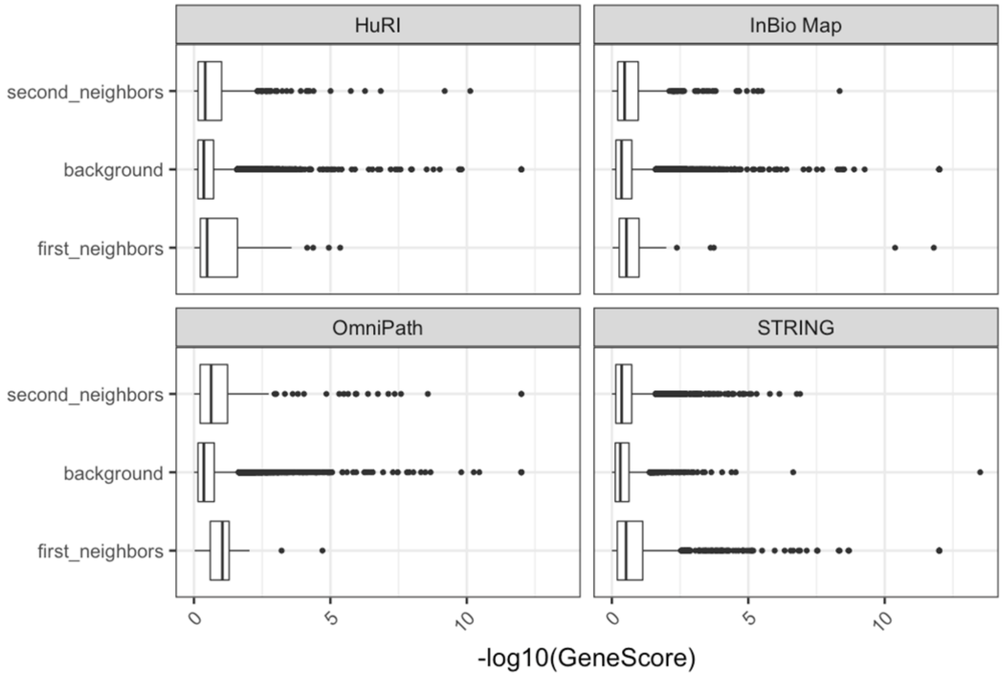


Supplementary Figure 4: A comparison of the Pascal gene score across the 4 networks for first and second neighbors of HCGHs.

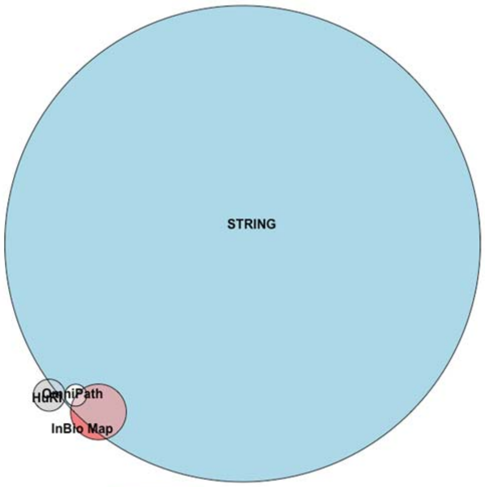


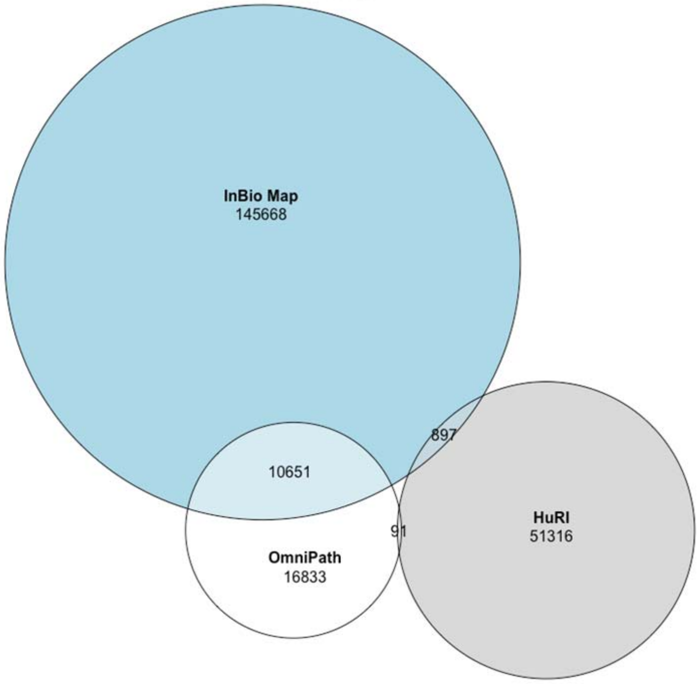


Supplementary Figure 5: The overlap in unique interactions across the different network resources used.

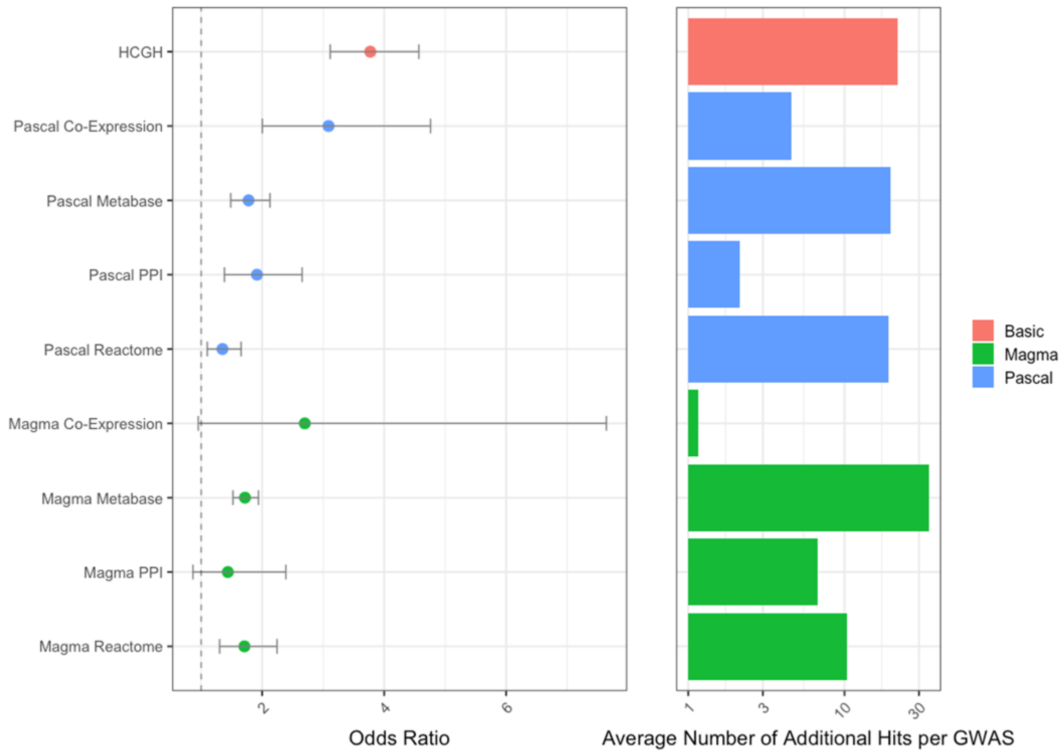


Supplementary Figure 6: Enrichment of successful drug targets (left) and number of targets implicated (right) for Pascal and Magma (with the HCGH reference)
